# Discovery of Primary, Cofactor, and Novel Transcription Factor Binding Site Motifs by Recursive, Thresholded Entropy Minimization

**DOI:** 10.1101/042853

**Authors:** Ruipeng Lu, Eliseos Mucaki, Peter K. Rogan

## Abstract

Data from ChIP-seq experiments can determine the genome-wide binding specificities of transcription factors (TFs) and other regulatory proteins. In the present study, we analyzed 745 ENCODE ChIP-seq peak datasets of 189 human TFs with a novel motif discovery method that is based on recursive, thresholded entropy minimization. This method is able to distinguish correct information models from noisy motifs, quantify the strengths of individual sites based on affinity, and detect adjacent cofactor binding sites that coordinate with primary TFs. We derived homogeneous and bipartite information models for 89 sequence-specific TFs, which enabled discovery of 24 cofactor motifs for 118 TFs, and revealed 6 high-confidence novel motifs. The reliability and accuracy of these models were determined via three independent quality control criteria, including the detection of experimentally proven binding sites, comparison with previously published motifs and statistical analyses. We also predict previously unreported TF cobinding interactions, and new components of known TF complexes. Because they are based on information theory, the derived models constitute a powerful tool for detecting and predicting the effects of variants in known binding sites, and predicting previously unrecognized binding sites and target genes.

## INTRODUCTION

Transcription factors positively or negatively interact with the regulatory elements in genes to mediate elaborate regulation of tissue‐ and stage-specific expression, which is a critical contributor to the marvelous complexity of the human body (1, 2). They could be divided into two general classes: one consisting of those TFs that are able to directly bind to DNA by recognizing specific sequence motifs, and the other comprising those TFs which indirectly bind to DNA by acting as interacting partners of the sequence-specific TFs (3). Some sequence-specific TFs have homogeneous binding motifs due to their independent binding activity, while others need to first form homodimers or heterodimers so that their binding sites are bipartite (4).

Not only do genome-wide interactions occur frequently between the two classes of TFs, but also abound between the sequence-specific TFs, with each interacting party being called a cofactor. For instance, NF-Y extensively coassociates with FOS over all chromatin states, cluster classes, and genic features, with 45% of NF-Y sites directly overlapping a FOS site and 39% of FOS sites directly overlapping an NF-Y site in K562 cells (5). NF-Y and FOS sites are located just as close (<50bp) as that observed between the NF-Y subunits or between FOS and JUN. CTCF extensively colocalizes with cohesins consisting of SMC1/SMC3 heterodimers and two non-SMC subunits RAD21 and SCC3 (6)

Although generally the binding sequences of TFs are highly conserved, some degree of variability at most positions of their binding sites is still evident in most TFs. Information theory-based models can be used to quantitatively and accurately describe base preferences in these motifs. A homogenous model for a TF is a positional weight matrix (PWM) can be derived from a set of aligned binding sites using Shannon information theory (7). A bipartite model consists of two PWMs, each of which corresponds to a half site, separated by a range of sequence gaps. Our laboratory previously developed software to generate information-theory based models of bipartite (and homogeneous) binding sites (4). The individual information content (*R*_*i*_) of a sequence bound by the TF, which represents the affinity of the TF-DNA binding site interaction, can be determined from these models. The *R*_sequence_ value of a model is the mean of the *R*_*i*_ values of all the binding site sequences used to compute the model, and represents the average binding affinity (8). ChIP-seq assays, which use chromatin immunoprecipitation (ChIP) to identify the repertoire of binding site sequences in a genome, can provide substrates for development of robust information models of TFs, and likely include binding site sequences of coregulatory TFs and potentially, cofactors.

Previous work related to the derivation of TF binding motifs has been based on experimental evidence supported by computational approaches. Weirauch et al. (9) made use of the protein binding microarrays (PBM) technique to determine TF sequence specificity for >1,000 TFs encompassing 54 different DNA binding domain (DBD) classes from 131 diverse eukaryotes. Jolma et al. (10) obtained 830 binding profiles representing 411 human and mouse TFs using high-throughput SELEX and ChIP sequencing. However, the oligonucleotide-based approach is not able to well account for variable-length spacers in bipartite binding sites, and it may produce multiple potential models among which correct ones, if present, may not be discriminated. In addition, the dynamic range of oligonucleotides used in the DNA microassays is limited so that not all the possible binding site sequences in the genome are covered as in the ChIP-seq assays, and there is no way to discover cofactors. Wang et al. (3) carried out de novo motif discovery for 119 human TFs from 457 ChIP-seq datasets using the MEME-ChIP software suite, and Kheradpour et al. (11) provided a systematic motif analysis for 427 human ChIP-seq datasets of 123 TFs using five motif discovery tools. However, these studies did not generate bipartite motifs with half sites separated gaps varying in length; more importantly, the derived motifs were only based upon strongest ChIP-seq signal peaks (top 500 or 250 peaks), effectively eliminating thousands of intermediate or weak binding events and biasing the resulting PWMs toward high-affinity, consensus-like binding sites. This is necessary, as the weakest binding sites inferred from ChIP-seq are essentially noise that can obfuscate the detection of true binding motifs. Nevertheless, this type of bias distorts the binding strengths estimated from the population of sites (12).

In this study, we developed a novel motif discovery approach that integrates recursive and thresholded features into the entropy minimization algorithm (4). We then applied this method to 745 ENCODE (Encyclopedia of DNA Elements [13]) ChIP-seq datasets of 189 human TFs. The majority of the primary binding motifs were successfully recovered and refined for the immunoprecipitated TFs, in addition to motifs for many known and previously unreported cofactors.

## MATERIALS AND METHODS

### ENCODE ChIP-seq datasets

The ENCODE Consortium conducted 907 ChIP-seq assays for human TFs and generated initial peak datasets for each replicate of each assay using the uniform peak calling pipeline (14), and for some assays further produced the optimal and conservative IDR thresholded peaks after applying the IDR (Irreproducible Discovery Rate) framework to the initial datasets to improve irreproducibility between multiple replicates of one assay. In addition, Factorbook (3,15) also provided some refined datasets directly generated by the SPP peak calling software (16).

We primarily used the IDR thresholded peak datasets, because we found that these data is more likely to discover primary or cofactor motifs than from the initial (i.e., unprocessed) datasets; and they contain more peaks and thus more binding sites than the truncated corresponding Factorbook datasets. One notable exception, however, was BRG1 which was found to exhibit motifs of cofactors GATA1 and AP1 exclusively from Factorbook, because IDR thresholded data produced only noisy low information content motifs. The initial datasets were examined if neither IDR-thresholded nor Factorbook datasets were available. We also noticed, that in some instances, that different technical replicates of the same ChIP-seq assay were not very consistent. For example, for one of the TEAD4 datasets, the motif for the NRSF TF was discovered from the initial peaks of replicate 1 on the A549 cell line, whereas both the cofactor AP1 and the primary motif were derived from replicate 2.

### Sequence motif discovery

Initially, information models from ChIP-seq reads were derived by entropy minimization with Bipad (5). However, we noticed that these models would not always contain the expected primary motifs and might show motifs of known cofactors of the primary TF. Additionally, datasets with an excess of weak and false positive peaks were noted to result in models consisting of overrepresented, noisy motifs. In order to enhance its ability to detect primary motifs in large datasets, we introduce a novel motif discovery algorithm by adding recursive masking and thresholding functionalities to entropy minimization (Figure 1). Multiple ChIP-seq datasets from distinct cell lines for the same TF, if available, were all examined for enriched sequence motifs to assess whether this approach was reproducible, and discover tissue specific preferences between these data sources.

**Figure 1.**
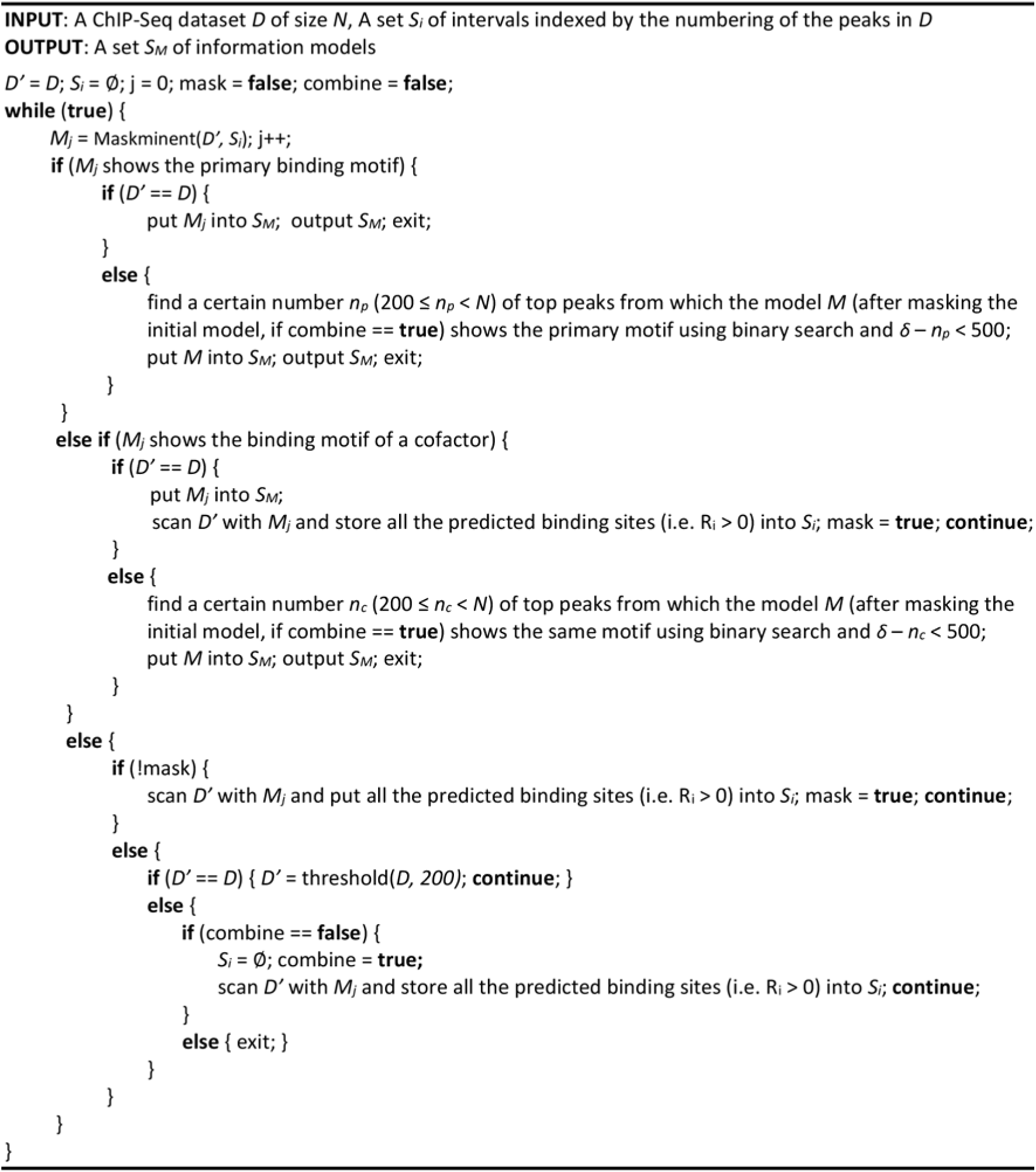
The motif discovery algorithm. Pseudocode describing the conversion of ChIP-seq data sets into information models using Bipad, a program which discovers sequence elements in unaligned sequences.

This masking technique provided a means of discovering additional conserved motifs in the same data by masking the motifs found in previous iterations of this modeling procedure. It is implemented by scanning a dataset with previous models and recording the coordinates of all the predicted binding sites, and then skipping these intervals when reanalyzing the dataset with Bipad. This is done to reduce the entire multiple alignment search space by the part covered by all the predicted sites. We rename and revise the objective function for building bipartite models to:

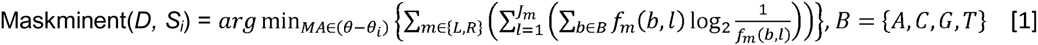

where *D* is the ChIP-seq dataset, *S*_*i*_ is the set of intervals of all the predicted sites of previous models to be masked, MA is one multiple alignment, *θ* is the multiple alignment space formed by all the peaks in *D*, *θ*_*i*_, is the search space covered by these predicted sites, *L* and *R* are the left and right half sites, *J* is the length of one half site, *f*_*m*_(*b,l*) is the frequency of the base *x* appearing at the position *l* of the half site *m*.

Thresholding the datasets to eliminate peaks with low signal strengths is another technique that can be used to reveal significant motifs distinguishable from noisy signals. Suppose that we want to take the top *n* peaks sorted by signal strength in a ChIP-seq dataset *D. size*(*D*) is the number of peaks in *D*, and *min*(*D*) and *max*(*D*) are the lowest and highest signal values among all the peaks in *D*, respectively. Then, the operation of thresholding the dataset *D* to obtain the top *n* peaks is defined by:

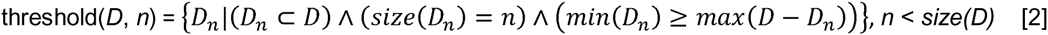

If the primary motif or a cofactor is unveiled by this operation, then theoretically there exists an accurate threshold *δ* on the number of top ranked peaks between showing the desired motif and not showing it and we will narrow down the error range within which *δ* is to less than 500 peaks. The masking and thresholding techniques can be combined to discover primary motifs.

Models are derived from ChIP-seq data with the Maskminent program, which implements the above algorithm. The ‐m option indicates whether to switch on the masking, otherwise the commands specifying the run to build homogeneous or bipartite models are similar to Bipad, respectively:

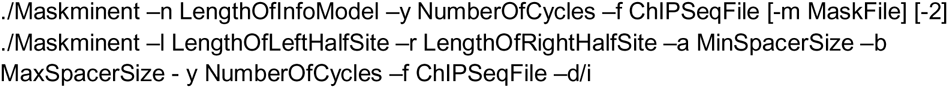

The Scan program is used to detect potential binding sites with an information model in a DNA sequence fragment based on the following parameters:

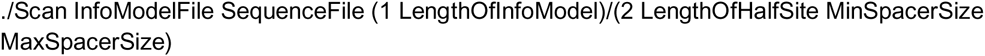

### Quality control

The quality control criteria used to evaluate the accuracy of our information models include:

1. Validating our models with experimentally proven binding sites in known target genes;
2. Comparing our models with the annotated motifs in the CIS-BP database (9) by determining their normalized Euclidean distances;
3. Evaluating the linearity of *R*_*i*_, values versus binding energy by performing F-tests on linear models and nested constant models between the two properties of predicted binding sites in our information models, in order to distinguish between correct and noisy, low complexity motifs.

We found that the *R*_*i*_, values are inversely proportionate to binding energy. The frequencies of binding sites appearing in a ChIP-seq dataset are linearly related to their binding energy (log_2_K_d_), which is delineated by Equation 1 derived by us starting from (17). And the *R*_*i*_, values of binding sites are proportionate to their frequencies, since the higher the affinity of a binding site is, the more the number of times it is bound by the TF will be in a ChIP-seq assay.

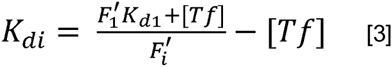

where *K*_*di*_is the dissociation constant of the *i*th predicted binding site, *F*_*i*_’ is its frequency, *F*_*1*_’ and *K*_*d1*_ are these values for the strongest site (i.e., the consensus sequence), and [*TF*] is the concentration of unbound TF. [*TF*] is negligible for most TFs.

The frequency of each binding site sequence within the multiple alignment of a model occurring in the ChIP-seq dataset was calculated as follows. If only one binding site occurred in a peak, its frequency would increase by one. In the case that a peak contained multiple binding sites, the frequency of each site would gain a fraction of one based on the proportion of its *R*_*i*_ value accounting for the sum of *R*_*i*_ values of all the sites. Then we applied Equation 3 to compute the binding energy of each site. The default value 10^−7^M of the dissociation constant *K*_*d1*_ was used, in instances where the exact value for a particular TF could not be established from the published studies.

## RESULTS

The motif discovery pipeline was applied to 745 ChIP-seq peak datasets of 189 TFs. The derived information models displayed primary motifs for 89 TFs (Supplementary Table S1 and S2), as well as 24 cofactor motifs for 118 primary TFs (Supplementary Table S3). We also describe 6 highconfidence novel motifs that have not been previously annotated.

### Primary binding motifs

*Homogeneous information models.* Correct information models were successfully derived for 62 TFs with homogenous binding motifs, which are concordant with published descriptions of these motifs (3).

Consensus or predominant sequences can present a distorted image of TF binding by missing sites of weak or intermediate strength (12). We determined the frequency of these most predominant sequences appear on a genome scale for 10 TFs (Figure 2A) by counting the peaks containing these sequences in their respective datasets. Surprisingly, only 0.015%-7.3% of all peaks contain binding sites with the most predominant sequence, demonstrating that these predominant binding sites are extremely rare in the ChIP-seq datasets. This is consistent with the observation that intermediate and low-affinity TF-DNA interactions are the most prevalent and abundant *in vivo* and can effectively regulate gene expression (18). In addition, models for NRSF clearly demonstrate that the core positions in its binding sites contain the predominant sequence motif, CTGTCC.

**Figure 2.**
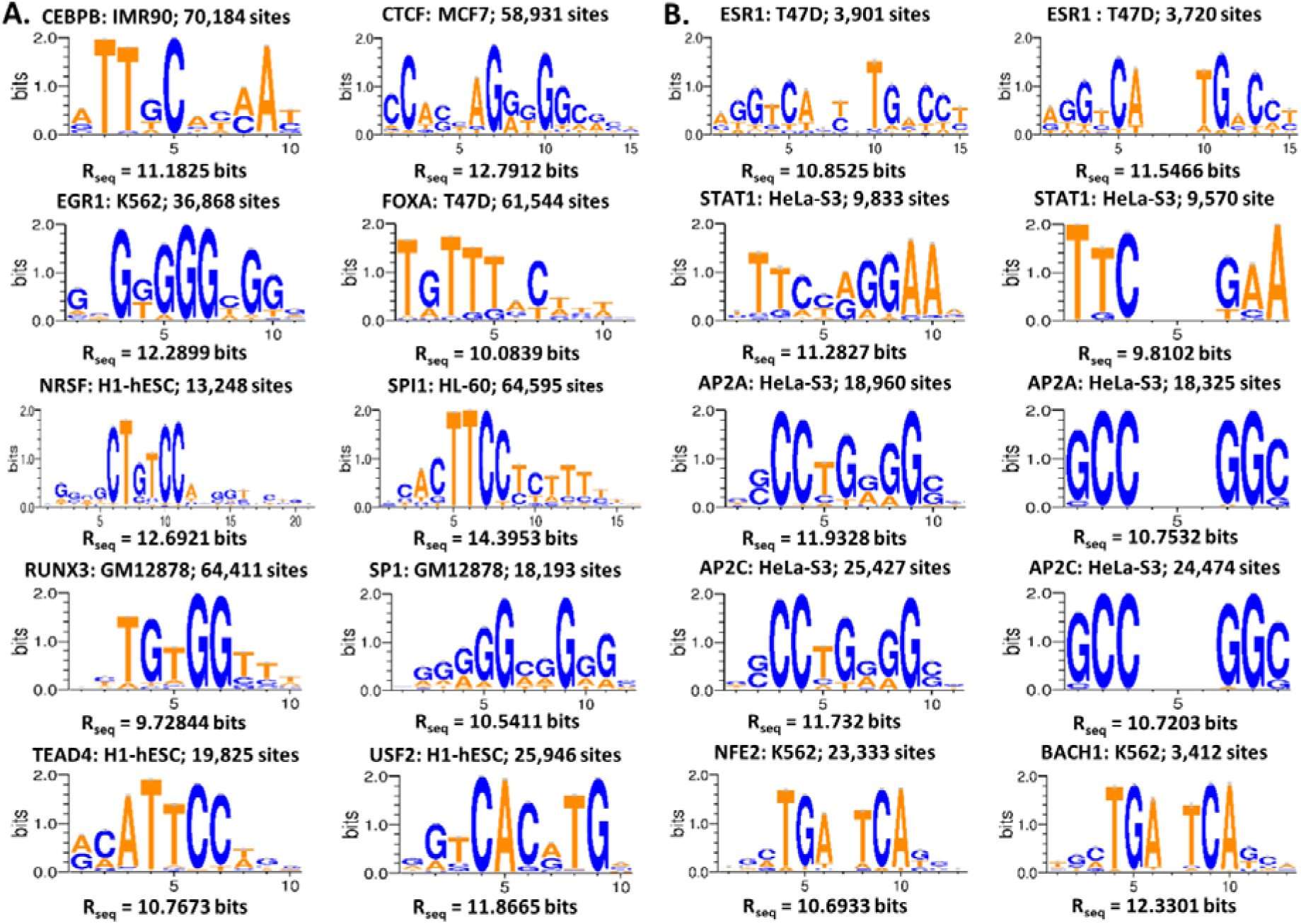
Sequence logos of homogeneous (A) and bipartite (B) information models. The TF name, and the cell line from which the model was derived, and the number of binding sites that the model contains are displayed with each logo. In (B), each of the first four rows includes a homogeneous (left) model and a bipartite (right) model of one TF from the same dataset. The last row includes the bipartite models of NFE2 (left) and BACH1 (right). The bipartite search patterns are 6<0,5>6, 3<2,4>3, 3<2,4>3, 3<2,4>3, 6<1,2>6 and 6<1,2>6, respectively from top to bottom.

### Bipartite information models

For 18 TFs bipartite information models were successfully derived, and were in agreement with previously reported motifs. The following demonstrates the validity of these models:

El Marzouk et al. (19) demonstrated that ESR1 is able to recognize binding sites of spacer lengths from 0bp to 4bp, with sites containing a 3bp spacer exhibiting the highest affinity. We allowed the spacer length to vary between 0 - 5bp from T47D cell line ChIP-seq data. Consistent with their findings, the resultant models show the documented predominant motif and are palindromic (Figure 2B). The bipartite model contains greater information contents than the corresponding homogeneous site, and the dominant gap size remains 3bp. However, the models contain some bipartite sites with positive *R*_*i*_ values and a 5bp gap, implying that ESR1 may be capable of binding to sites that were not previously examined. The symmetry between the half sites exhibited by the bipartite models suggests that dimeric ESR1 may bind a narrow range of sequences with similar half site affinities.

The palindromic predominant motif of the AP2 family is 5’-GCCN_3_GGC-3’, and other binding sequences confirmed in an *in vitro* binding-site selection assay include 5’-GCCN_4_GGC-3’ and 5’-GCCN_3/4_GGG-3’. Another binding site 5’-CCCCAGGC-3’ was also found in the SV40 enhancer (20). Indeed, the gaps in the bipartite models of AP2A and AP2C range from 2bp to 4bp, which is concordant with published reports; therefore, these models are representative of the genome-wide pool of *bona fide* binding sites (Figure 2B). Besides the fact that they are quite symmetric, we also observed that the two outermost positions allow the highest degree of variability and that adenine can also appear at the first position of the right half site apart from the consensus guanine base. The bipartite models of AP2A and AP2C exhibit the same pattern of conservation levels across all the individual positions, indicating that the two members of the AP2 family are almost indistinguishable in binding specificity.

STAT1 binding was reported to have a major gap size of 3bp and be able to bind to sites with a 2bp gap (N_2_), but appear to be incapable of recognizing N_4_ sites (21). However, our bipartite models for STAT 1 include a small number of binding sites with a 4bp gap, compared to those with 3bp or 2bp spacers (Figure 2B), which suggests that N_4_ sites can be recognized. The left‐ and rightmost nucleotides of each half site are nearly invariant, whereas the inner 4 nucleotide contacts are vary significantly.

NFE2 and BACH1 form heterodimers with small MAF family proteins (MAFF, MAFG and MAFK) which recognize two types of bipartite palindromic motifs defined by the predominant binding sequences, TGCTGA(C)TCAGCA and TGCTGA(CG)TCAGCA (22). The previously reported binding motifs (3) are homogeneous, and do not account for the dimeric interaction that gives rise to the bipartite motifs. The bipartite models that we derived (Figure 2B) demonstrate that the inner six positions surrounding the dominant 1bp gap are more highly conserved than the outer six positions in the binding site.

*Discrepancies between information models of the same TF in distinct cell lines.* In several instances, we noticed a potential discrepancy between models of the same TF from different cell types, in both homogeneous and bipartite motifs.

For instance, among the three homogeneous models of ESR1 from ECC1 cells one half site has very high conservation levels and the information content of the other half site is lower, while the two half sites are much more symmetric in the T47D models (Figure 3A). For MAFF and MAFK, the discrepancy between the symmetric bipartite models from K562 and HepG2 cells is prominent; the outer 6 positions of these HepG2 models show a greater degree of conservation than the internal 6 positions, but the K562 models illustrates the opposite trend (Figure 3A). Additionally, MAFK motif derived from IMR90 cells resembles the HepG2 model, while the models from HeLa-S3 and H1-hESC datasets resemble the K562 model. This could be accounted for by differences in the compositions of binding sites, i.e., different target genes for the same TF in different tissues.

**Figure 3.**
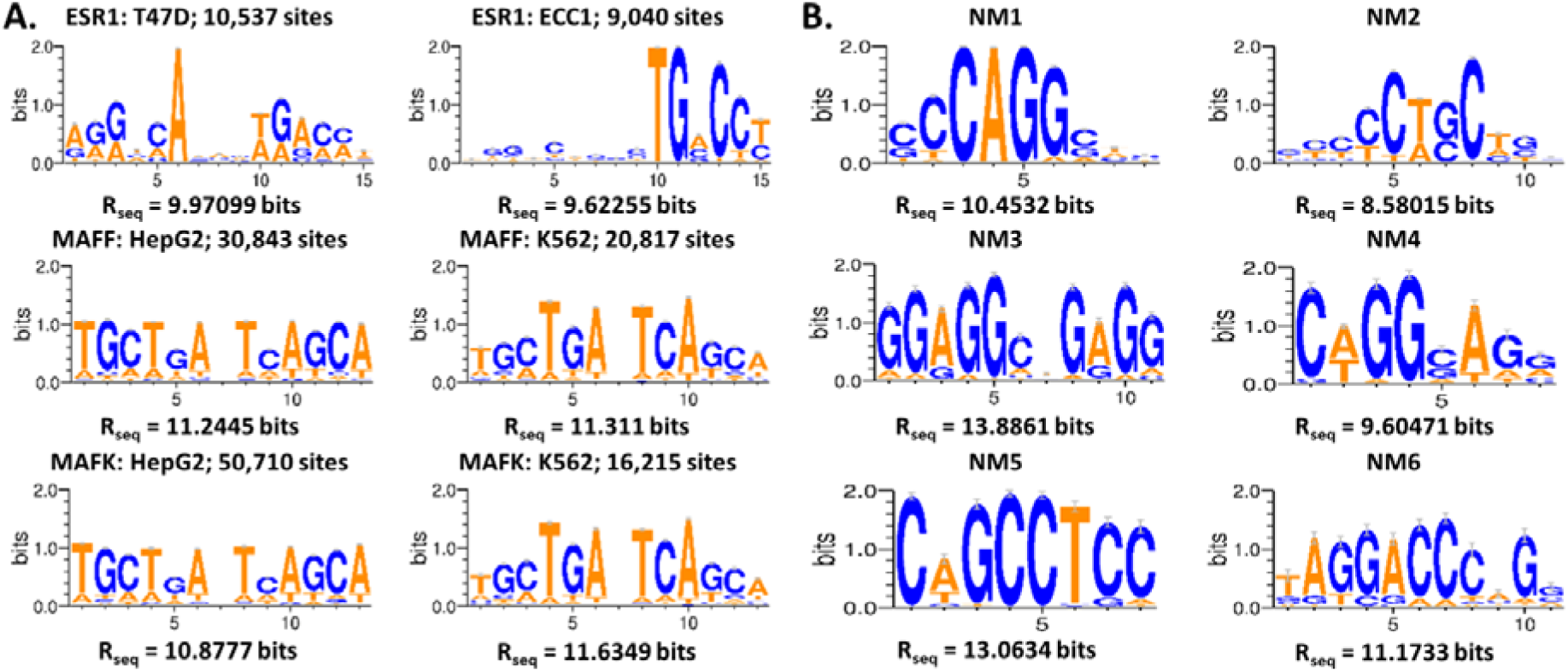
Comparison between information theory-based models from different cell lines and motifs. (A) Each row includes sequence logos of two models of the same TF from two different cell lines. Note: the bipartite models for MAFF and MAFK used the search pattern 6<1,2>6. (B) The high-confidence novel motifs (“NM1” –“NM6”).

### Cofactor binding motifs

Discovery of the binding motif of a cofactor in one dataset of a particular primary TF suggests that there exists transcriptional co-regulation between the two TFs on their common target genes, either by binding of the cofactor in the complex formed by their physical interactions, or by their cobinding to achieve a synergistic *cis*-regulatory effect. *De novo* motif discovery from ChIP-seq datasets provides an effective approach for confirming or predicting the genome-wide TF interactions, in contrast to the abundant yet scattered existing literature evidences in specific genes about such interactions. Figure 4 illustrates the cofactors and their interactions with the primary TFs revealed by our motif discovery algorithm.

**Figure 4.**
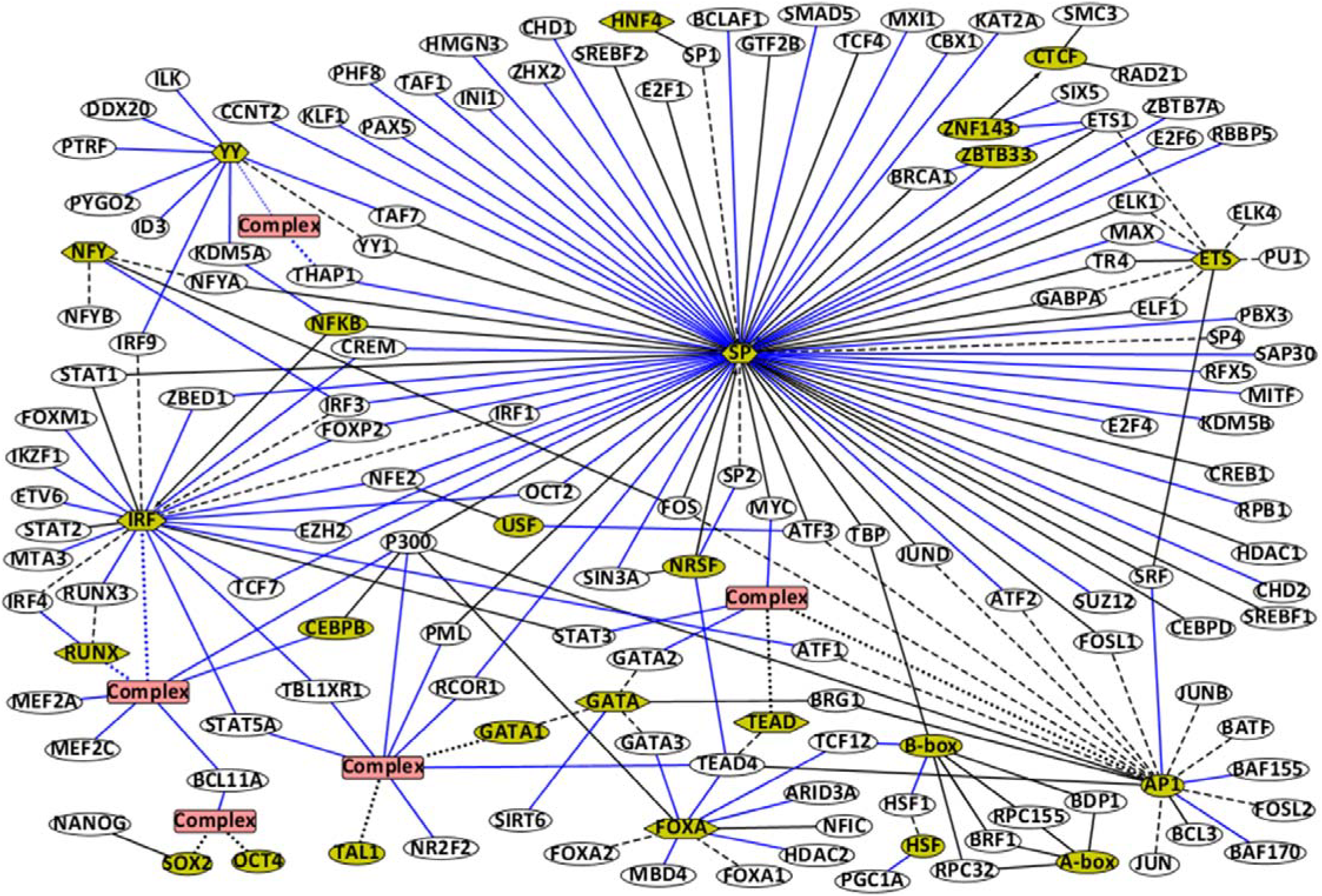
Network graph of cofactors and their interactions with the primary TFs revealed by our motif discovery pipeline. A yellow ellipse denotes a cofactor and a white ellipse denotes a primary TF. A hexagon denotes a TF family with dash lines connecting its members. For a TF family only members for which ENCODE provides peak datasets are shown. A red rectangle denotes a known or predicted complex with black or blue dotted lines indicating its components, respectively. An undirected line denotes the interaction between a primary TF and a cofactor which may be a complex or a TF family. A directed line links two cofactors, denoting that in a dataset of the starting TF the ending TF was discovered as a cofactor. Black lines denote known interactions and blue lines denote the newly discovered interactions.

#### Confirmation of known cofactors

Our information models confirmed genome-wide interactions between 15 cofactors and 50 primary TFs, for which we were able to find specific evidences in literature (5, 6, 23–65). For example, the interaction between SP1 and multiple members of the ETS and AP1 families is well characterized (66–71). ELK1 and SRF can recruit each other to form a ternary complex on the CArG-ETS elements (72). TEAD-AP1 cooperation with SRC coactivators serving as a bridge drives downstream gene transcription to regulate cancer cell migration and invasion (73). And STAT1, STAT2 and IRF9 can form a heterotrimer that regulates transcription of genes containing IFN-stimulated response elements (ISREs) (74). The GATA1-TAL1 and SOX2-OCT4 complexes emerged from the datasets of TAL1 and OCT4 as primary motifs, respectively, suggesting the formation of the two complexes is necessary for the functioning of TAL1 and OCT4.

#### Discovery of novel cofactors

A number of previously unrecognized, probable cofactor motifs (n=19) were revealed by our algorithm for 87 primary TFs. Enrichment of binding sites of cofactors in the primary TF datasets provides evidence for these TF cobinding and interactions, and new subunits of known complexes. This includes possible associations between the IRF and RUNX families, and their further cooperation with BCL11A, MEF2A, MEF2C, CEBPB and P300 in GM12878 cells. Another similar prediction is that the TEAD-AP1 complex can recruit MYC, STAT3 and GATA2 in multiple cell lines. We also predict the existence of an YY1-THAP1 complex based on the co-emergence of their binding motifs in the K562 dataset of THAP1. Other predictions include coordination between USF and ATF3 across multiple cell lines from different tissues, between ZNF143 and SIX5 in multiple cell lines, between ZNF143 and ETS1 in A549 and K562 cells, between ZBTB33 and ETS1 in GM12878 lymphoblastoid cells, between the FOXA family and GATA3 in T47D cells, between FOXA and ARID3A, TCF12, TEAD4 in HepG2 cells, between AP1 and SRF in HCT116 cells, between the YY family and IRF9 in K562 cells, between NFY and IRF3 in HeLa-S3 and HepG2 cells, between MAX and the ETS family in NB4 cells, between NRSF and TEAD4 in A549 cells, between NRSF and SP2 in HepG2 cells, between the SOX2-OCT4 complex and BCL11A in H1-hESC cells, between the TFIIIC complex and HSF1 in HepG2 cells, between TFIIIC and TCF12 in MCF-7 cells, between the IRF family and 11 DNA-binding TFs in GM12878 and K562 cells, and between the SP family and 23 DNA-binding TFs in multiple tissues. A DNA-binding complex consisting of GATA1, TAL1, E2A, LMO2 and LDB1 is present in the erythroid cell lineage (40). Based on the proximity and coprecipitation of these binding sequences, suggest that this complex in which GATA1 and TAL1 contact DNA can coordinately bind with NR2F2, STAT5A, and TEAD4. The complex may also incorporate P300, PML, RCOR1 and TBL1XR1, which are non-DNA binding proteins.

Many cofactors were also discovered in datasets of non-sequence specific TFs, which likely means that these primary TFs are recruited to genes by associating with the DNA-binding cofactors. Our predictions based on this include interactions between NFKB and KDM5A in H1-hESC cells, between FOXA and HDAC2, MBD4 in HepG2 cells, between ZBTB33 and BRCA1 across multiple cell lines, between GATA and SIRT6 in K562 cells, between IRF and MTA3, TBL1XR1 in GM12878 cells, between IRF and EZH2 in GM12878 and K562 cells, between AP1 and SMARCC1, SMARCC2 in HeLa-S3 cells, between HSF1 and PGC1A in HepG2 cells, between YY and KDM5A in H1-hESC cells, between YY and DDX20, ID3, ILK, PTRF, PYGO2, TAF7 in K562 cells, and between SP and 18 non-sequence specific TFs in multiple tissues.

To validate the predicted cobinding between cofactors and primary TFs, we investigated distributions of their intersite distances by scanning the ChIP-seq datasets with our information models (Figure 5, Supplementary Table S4). We applied a threshold (e.g. R_sequence_ or ½ * R_sequence_ if using R_sequence_ leads to few binding sites of cofactors) on the *R*_*i*_ values of predicted binding sites in order to remove the relatively large number of weak binding sites that are likely to be low-complexity sequences. The SOX2-OCT4 complex was used as a primary negative control, as it is merely expressed in the H1-hESC cell line and thus cannot be a cofactor for the primary TFs in other cell lines. It can be seen that the percentage of peaks in which intersite distances between the discovered cofactor and the primary TF are quite small (e.g., less than 20bp) is overwhelming in the respective dataset, while there is no such a trend for the negative control and the primary TF. The distribution of intersite distances between TEAD4 and AP1, whose extensive colocalization was already observed in literature as mentioned above, was also generated. There exists the same difference between it and the distribution for TEAD4 and the negative control.

**Figure 5.**
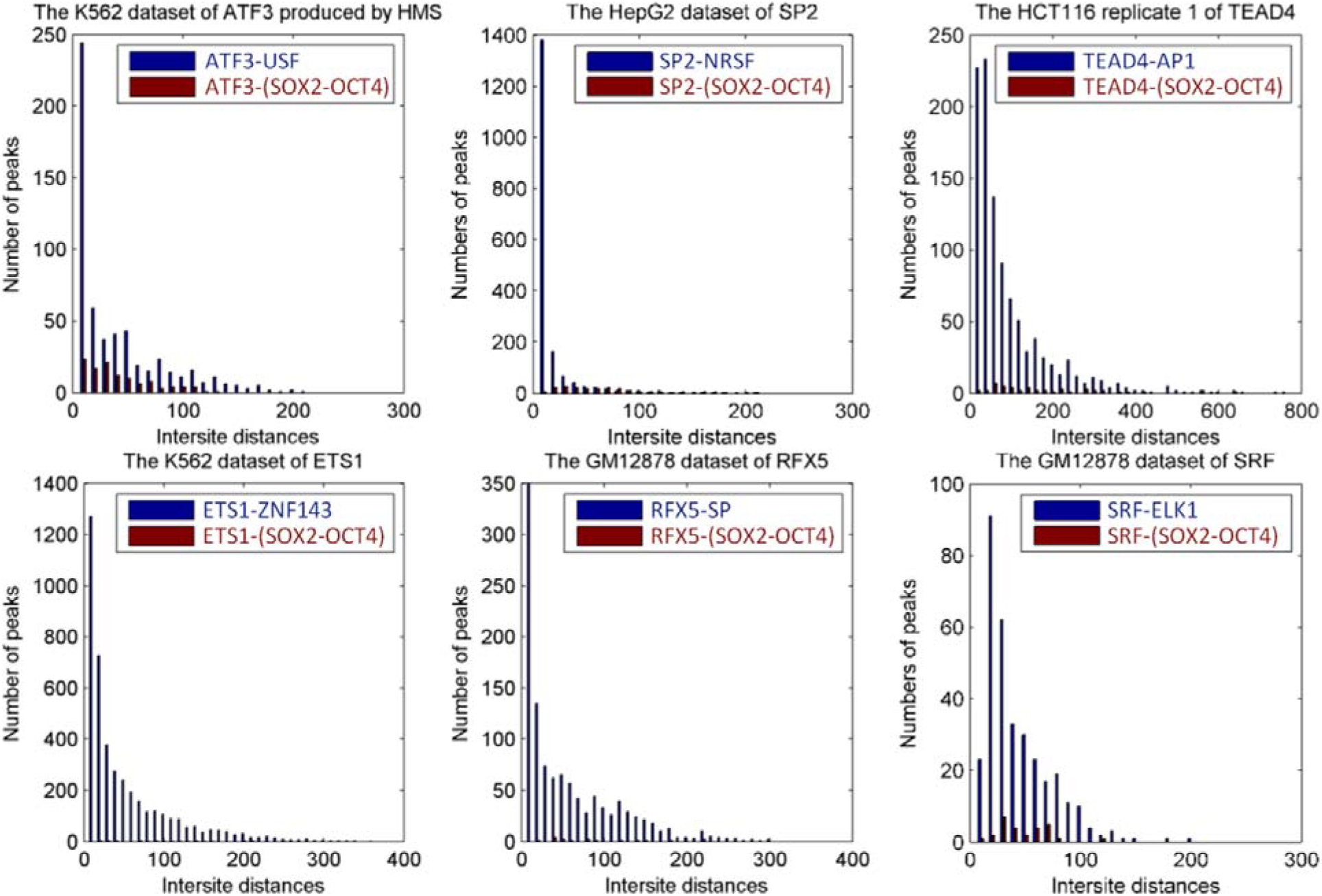
Distributions of intersite distances between primary TFs and discovered cofactors versus negative controls. The minimum threshold information contents of predicted binding sites is R_sequence_, except for ATF3, SP2 and TCF12, where it was set to 0.5 * R_sequence_ if using R_sequence_ leads to few binding sites of cofactors. Each graph illustrates a much higher frequency of short (< 20nt) intersite distances between primary TFs and cofactors (blue) compared to the negative control (SOX2-OCT4; red).

Further detailed investigation into the binding sites of cofactors and primary TFs in peak datasets revealed that a lot of binding sites of the IRF and RUNX families overlap, which can be understood by the common T-rich feature of their predominant sequences NNTTTCNNTTTC and TGTGGTTT. This phenomenon was also observed for AP1 and TEAD4. The predominant sequences of AP1 and TEAD4 are TGA(‐/‐ ‐)TCA and (A/G)CATTCC(T/A)G(G/C), and the prevalent overlapping pattern is that the left half site TGA of an AP1 binding site is at the same positions with the rightmost three bases of a TEAD4 site, or the right half site TCA is shared by the leftmost three bases of a TEAD4 site. Similarly, a large quantity of AP1 and USF binding sites intersect. Given the USF predominant sequence is CA(C/T)GTGAC, the common intersecting pattern is that the first two bases TG of an AP1 site are shared by the last two bases of the E-box in a USF site, and vice versa.

*Tissue-preferential activity of cofactors and their tissue-specific distributions relative to primary TFs.* Several cofactors were revealed recurrently with different primary TF partners, and notably in specific cell lines. This is consistent with the possibility of coordinate regulation with these primary TFs in a tissue-preferential manner. For example, all the datasets of 24 primary TFs in which the IRF family was discovered as a cofactor came from GM12878 or related cell lines, except that the ones of STAT1, STAT2, ATF1 and one dataset of EZH2 were from K562 cells, and the one of FOXP2 was from SK-N-MC cells. The GM12878 cells are of B-lymphoblastic type, which is concordant with the critical role that the IRF members play in the regulation of the immune system (75). All the datasets of 10 primary TFs from which the GATA and GATA1-TAL1 motifs emerged as cofactors were derived from K562 cells of the erythrocytic leukemia type, which is supported by the role the GATA family exhibits in hematopoietic lineage gene expression, e.g., GATA1 being an erythroid-specific activator (76, 77). Similarly, FOXA was revealed to be a cooperating partner in datasets of 8 primary TFs from the HepG2 cell line derived from hepatocellular carcinoma cells, except that two datasets of GATA3 and P300 were produced from the T47D cell line representing breast cancer cells. This suggests that the FOXA proteins, which are known to regulate the initiation of liver development (78), are highly active in liver tissue. Another TF family known to be a key factor regulating hepatocyte differentiation and liver-specific functions is HNF4 (79), which was discovered as a cofactor of SP1 in a HepG2 dataset. SOX2 and the SOX2-OCT4 complex were unveiled as cofactors only in datasets of 3 primary TFs from the H1-hESC cell line representing embryonic stem cells (ESC), which is in agreement with the fact that SOX2, OCT4 and NANOG are essential for maintaining ESC pluripotency (80). Interestingly, all the datasets (n=9) in which YY was revealed as a cofactor were from K562 cells, except that the one of KDM5A was from H1-hESC cells. Unlike GATA, the YY family is ubiquitously distributed and not known to play an especially central role in erythroid cells, although it was reported that YY1 acts as a developmental repressor of the erythroid-specific ε-globin gene along with GATA1 (81)

The SP family was found to be capable of interacting with the maximum number of TFs, which is consonant with its ubiquitous expression and may be related to its relatively simple purine-rich binding motif. Similarly, the ubiquitously expressed AP1 could interact with 10 TFs in multiple cell lines without showing an extreme cell type preference.

A number of primary TFs exhibit an extensive capability of interacting with multiple cofactors in different tissues. The unique distribution of these cofactors across multiple cell lines suggests the tissue-specific functions of the primary TFs. For instance, TEAD4 was found to be a cooperative partner of GATA1-TAL1 in K562 cells, of NRSF in A549 cells, of FOXA in HepG2 cells, and of AP1 in multiple tissues. Cofactors which were discovered to interact with P300 include IRF/RUNX in GM12878 cells, SP in H1-hESC cells, AP1 and CEBPB in HeLa-S3 cells, FOXA in HepG2 and T47D cells and GATA1-TAL1 in K562 cells. Based on cosegregation there is an interaction between BCL11A and IRF/RUNX in GM12878 cells, and SOX2-OCT4 in H1-hESC cells. STAT5A and TBL1XR1 cosegregated with members of the IRF family in GM12878 cells and with GATA1-TAL1 in K562 cells.

### Novel binding motifs

We uncovered 6 high-confidence novel motifs that have not been previously annotated (Figure 3B). The “NM1” motif was considerably enriched in the datasets of non-DNA-binding proteins, BAF155 and BRG1, from HeLa-S3 cells and the “NM2” motif was highly conserved in the datasets of BCL11A and NANOG from H1-hESC cells.

The “NM3” motif was revealed in the datasets of ESRRA and SREBF2 from GM12878 cells, MAX from HCT116 cells, CREB1 and GTF3C2 from K562 cells, non-DNA-binding RCOR1 from IMR90 cells. The Euclidean distances between these novel motifs and primary motifs range from 3.1 - 3.4 bits per nucleotide. The “NM4”, “NM5” and “NM6” motifs were discovered in the datasets of GATA3, MXI1 and FOSL1 from MCF-7, SK-N-SH, and H1-hESC cells, respectively. These distances range from 2.9 to 3.4 bits per nucleotide. These motifs may comprise the binding motifs of currently unknown TFs or other non-annotated functional elements *in vivo.*

### Quality control

*Validation of information models with bona fide binding sites.* We found a total number of 794 experimentally confirmed binding sites in literature for all of the 89 TFs whose primary binding motifs were discovered, and attempted to detect these sites using our information models, in order to assess the credibility and accuracy of our models well capturing the genuine binding specificities of these TFs. It turned out that our models did not fail to detect a single true binding site. The detailed results are listed in Supplementary Table S5.

*Comparison between our information models and other binding motifs.* Binding motifs of eukaryotic TFs were previously reconstructed from oligonucleotide binding selection assays (9); these motifs represent a type of ground truth reflecting the bona fide sequence preferences of these TFs. For 133 TFs, we compared our information matrices with these motifs quantitatively by determining the normalized Euclidean distances between them, and classified the distances into three categories (Table 1). We observed that, for most TFs, the models derived in this study and the reconstructed motifs are nearly identical or only differ at 1 or 2 positions. The discovery of cofactors by recursive, thresholded entropy minimization was the predominant explanation for large normalized distances for 39 of these TFs.

**Table 1:** Comparison of information-theory based models with the motifs derived by reconstruction of bound, overlapping oligonucleotides

| Normalized Euclidean<br>Distance (bits per nucleotide) | Implication | Number of TFs in Category |
| --- | --- | --- |
| <1 | Nearly identical models | 75 |
| 1-2 | Models differ at 1 – 2 positions | 18 |
| >2 | Usually cofactor; sometimes<br>noise | 39 |

#### Statistical analyses on information models

An F-test was carried out on linear regression models to assess the distributions of the *R*_*i*_ values and binding energy of putative binding sites in each information model. F-test values on 60 models showing primary binding motifs and 60 noise models randomly selected were calculated (Figure 6; data available in Supplementary Table S2). The F-test values of 70% correct models (42/60) are above 1,000, while all the noise models have F-test values below 1,000, making 1,000 an appropriate threshold to distinguish between correct and noise models.

**Figure 6.**
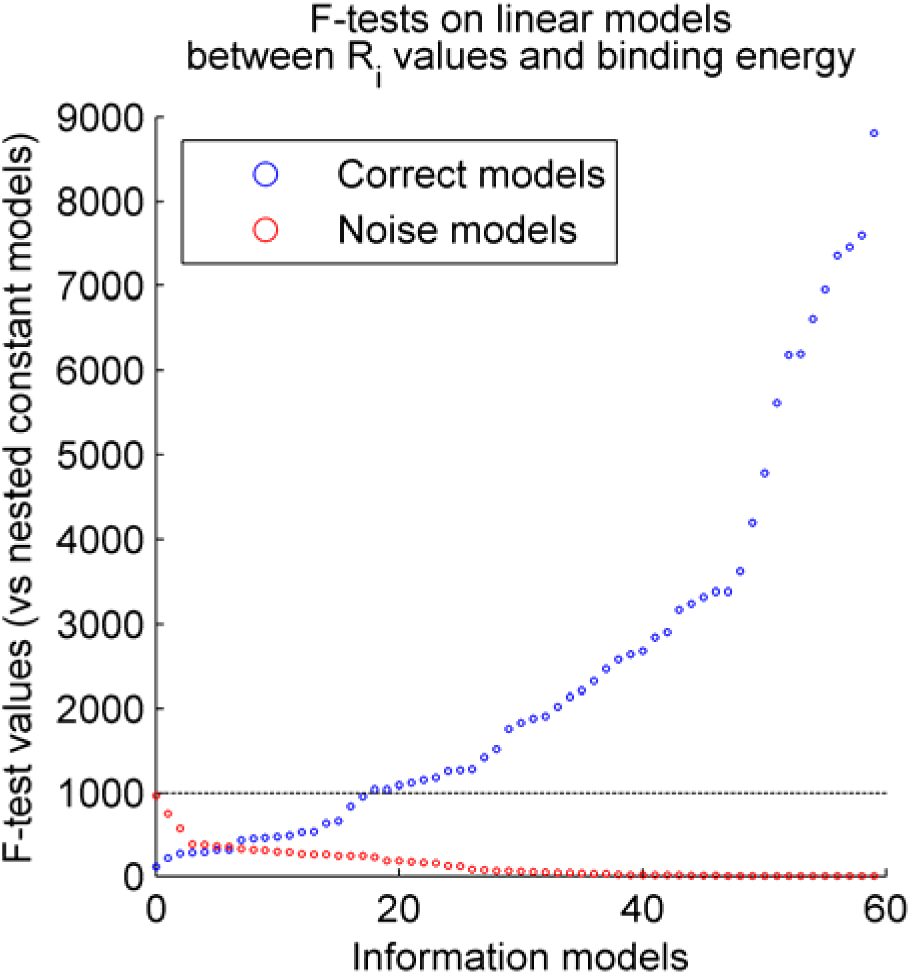
F-test results on linear models between *R*_*i*_, values and binding energy. F-tests performed on 60 models showing primary binding motifs (blue circles) and 60 noise models (red circles) show all noise models have F-values below 1000 while the F-values of the majority primary motifs are above 1000. Thus as indicated by the dash line, 1000 is an appropriate threshold for distinguishing correct from noise models.

## DISCUSSION

Using our motif discovery pipeline based on recursive, thresholded entropy minimization, from ENCODE ChIP-seq datasets of 189 human TFs we obtained accurate homogeneous and bipartite information models showing primary binding motifs for 62 and 18 TFs, respectively. In addition, we discovered 24 cofactor motifs for 118 TFs and 6 high-confidence novel motifs as well. Our findings lend support for predictions of known and novel TF-cofactor interactions, new subunits of known complexes, and can reveal target genes that previously, were not known to be regulated by both TF and one or more cofactors. In addition to homogeneous information models, bipartite models with variable-length gaps, which more precisely reflect the binding behavior of dimeric TFs, were also generated. Our approach is generally able to distinguish correct models from noisy, low complexity motifs that are coincidentally enriched in the ChIP-seq data. The models do not only take high-affinity binding sites into consideration, but also encompass many intermediate and lower-affinity sites. This level of accuracy justifies using these models to detect variants in known binding sites that alter the affinity of their interactions with their cognate TFs.

Within a peak from a ChIP-seq dataset, the strength of the binding site of the primary TF is expected to be proportionate to the integrated signal intensity of the peak. Therefore, the average strengths of the binding sites contained in a subset of top ranked peaks is greater than the remaining sites in a dataset; while this introduces some bias, this subset comprises many more sequences than those found in Factorbook for any of the transcription factors. Without this thresholding, the minimum entropy generates noisy or cofactor models, and therefore, top ranked peaks filtering more effectively reveals primary TF motifs. The masking and thresholding techniques decrease the multiple alignment search space from distinct directions, with masking removing a portion in which the initial model is contained, and thresholding increasing the possibility of a non-noise motif possessing the smallest entropy in the remaining search space.

In addition to PWMs, high-throughput DNase-seq assays have also been used to predict TF binding sites across the genome in literature, which is based on how similar a footprint shape of a candidate site is to an absolute DNase-seq footprint. In order to compare the ability of our information models and DNase-seq predicting direct binding, we obtained DNase-seq data on true binding sites of 20 TFs within ChIP-seq peaks from Yardimci et al. (82), and attempted to detect these sites using our models. All these binding sites were successfully detected by our models, whereas DNase-seq footprinting was only able to identify 35%-85% of them, when 0 was used as the threshold on FLR values indicating the similarity between digestion shapes of candidate sites and definite footprints (Supplementary Table S6). As weak binding sites tend not to generate footprints and thus not to be discovered by DNase-seq, the expectation is that the sites detected by DNase-seq would be stronger than those that evade detection. In fact, this trend was observed for only 10 TFs and the average strengths of these classes of these binding sites were not significantly different.

These information theory-based models not only are capable of detecting strong binding sites, but also discover intermediate and weak sites (Supplementary Table S4). This can be ascribed to the fact that as many peaks as possible were retained when deriving these models from ChIP-seq datasets, under the precondition that the correct binding motifs were shown. By including weaker binding sites in addition to those resembling consensus-like motifs, the weights in the R_i_(b,l) PWMs are more representative of the full range of binding affinities. The resultant *R*_*i*_ values are more effective for forecasting potential binding sites and target genes, detecting variants in known binding sites and predicting their effects. Our models can also be used to forecast whether a SNP (Single Nucleotide Polymorphism) linked to a disease risk lies within a TF binding site (83) as a means of modulating expression of these genes, and potentially offer new explanations for disease etiology in some individuals. The novel cofactor binding sites in TF gene targets suggest previously unrecognized coregulation mechanisms in which they may be operative in these genes.

In fact, the number of TFs for which cofactor motifs were revealed exceeds the number of TFs whose primary binding motifs were discovered, partially because only cofactor motifs can be found in the datasets of TFs which exhibit little or no sequence specificity (e.g., CCNT2, INI1 and P300). For 11 primary TFs, the binding site sequences were extremely variable, that is, the overall conservation levels of their binding motifs contain less information than noisy, low complexity sequences or cofactor motifs. Entropy minimization finds the highest information content motif in a ChIP-seq dataset for a given length motif and degree of sequence conservation over that length. For 18 primary TFs associated with cofactors, which themselves physically contact DNA, the primary TF motif was not enriched. The inability of the software to discover such primary motifs is a limitation of this approach. Interactions between the primary TFs and a subset of the cofactors which are known to cooperate with them were detected, since the association has to occur with a prevalence sufficient to produce a recognizable motif (usually >0.5 bits per nucleotide over the entire site). Nevertheless, the algorithm will miss some cofactors with weakly conserved or that overlap with other conserved motifs.

While unable to discover cofactors nor identify bipartite motifs of variable spacing, the PBM technique adopted by Weirauch et al. (9) and Jolma et al. (10) is able to determine binding specificities for a wide range of sequence-specific TFs, in that homogeneous binding sites of TFs are reconstructed from overlapping oligonucleotide sequences by directly detecting complexes with the TF. This eliminates interference of noisy sequences or cofactors which may emerge as false minimum entropies using our method. The configuration of outputting up to five significant motifs in MIME and the simultaneous use of five motif discovery tools lead Wang et al. (3) and Kheradpour et al. (11) to discovering a few cofactors that were not revealed by our method, albeit using only top 500 or 250 peaks increased the likelihood of those cofactors appearing by chance.

Our motif discovery approach can be applied to other ChIP-seq data, to derive information models of primary TFs and to reveal cofactors that complex with primary TFs. The quality control criteria we described are capable of ensuring that the user-built models are accurate and can be used for binding site detection. The first quality control criterion is particularly important, because it provides a straightforward assessment of model performance. The recursively thresholded feature is crucial for guaranteeing that the discovered cofactors do not appear by chance, because the greater the number of peaks from which a cofactor is derived, the higher the confidence that the cofactor indeed interacts with the primary factor.

Our study comprehensively investigated a new approach to define TF binding specificities based on the ChIP-seq TF data that ENCODE has released, and mined a wealth of valuable information about TF binding motifs and interactions on a genome scale. This information expands the granularity of the current knowledge on TF interaction with DNA and points out potential directions for future experimental study on interaction between TFs.

## SOFTWARE AVAILABILITY

http://dx.doi.org/10.5281/zenodo.15692

## ACKNOWLEDGEMENT

We thank SHARCNET and Compute Canada for providing high performance computing facilities.

## REFERENCES

1. Leung,K.K., Ng,L.J., Ho,K.K., Tam,P.P. and Cheah,K.S. (1998) Different cis-regulatory DNA elements mediate developmental stage‐ and tissue-specific expression of the human COL2A1 gene in transgenic mice. J. Cell Biol., 141, 1291–1300.

2. Levine,M. and Tjian,R. (2003) Transcription regulation and animal diversity. Nature, 424, 147–151.

3. Wang,J., Zhuang,J., Iyer,S., Lin,X., Whitfield,T.W., Greven,M.C., Pierce,B.G., Dong,X., Kundaje,A., Cheng,Y., et al. (2012) Sequence features and chromatin structure around the genomic regions bound by 119 human transcription factors. Genome Res., 22, 1798–1812.

4. Bi,C. and Rogan,P.K. (2004) Bipartite pattern discovery by entropy minimization-based multiple local alignment. Nucleic Acids Res., 32, 4979–4991.

5. Fleming,J.D., Pavesi,G., Benatti,P., Imbriano,C., Mantovani,R. and Struhl,K. (2013) NF-Y coassociates with FOS at promoters, enhancers, repetitive elements, and inactive chromatin regions, and is stereo-positioned with growth-controlling transcription factors. Genome Res., 23, 1195–1209.

6. Parelho,V., Hadjur,S., Spivakov,M., Leleu,M., Sauer,S., Gregson,H.C., Jarmuz,A., Canzonetta,C., Webster,Z., Nesterova,T., et al. (2008) Cohesins functionally associate with CTCF on mammalian chromosome arms. Cell, 132, 422–433.

7. Shannon,C.E. (1948) A Mathematical Theory of Communication. Bell Syst. Technol. J., 27, 379–423, 623–656.

8. Rogan,P.K., Faux,B.M. and Schneider,T.D. (1998) Information analysis of human splice site mutations. Hum. Mutat. 13(1), 82.

9. Weirauch,M.T., Yang,A., Albu,M., Cote,A.G., Montenegro-Montero,A., Drewe,P., Najafabadi,H.S., Lambert,S.A., Mann,I., Cook,K., et al. (2014) Determination and inference of eukaryotic transcription factor sequence specificity. Cell, 158, 1431–1443.

10. Jolma,A., Yan,J., Whitington,T., Toivonen,J., Nitta,K.R., Rastas,P., Morgunova,E., Enge,M., Taipale,M., Wei,G., et al. (2013) DNA-binding specificities of human transcription factors. Cell, 152, 327–339.

11. Kheradpour,P. and Kellis,M. (2014) Systematic discovery and characterization of regulatory motifs in ENCODE TF binding experiments. Nucleic Acids Res., 42, 2976–2987.

12. Schneider,T.D. (2002) Consensus sequence Zen. Appl. Bioinformatics, 1, 111–119.

13. ENCODE Project Consortium (2012) An integrated encyclopedia of DNA elements in the human genome. Nature, 489, 57–74.

14. Landt,S.G., Marinov,G.K., Kundaje,A., Kheradpour,P., Pauli,F., Batzoglou,S., Bernstein,B.E., Bickel,P., Brown,J.B., Cayting,P., et al. (2012) ChIP-seq guidelines and practices of the ENCODE and modENCODE consortia. Genome Res., 22, 1813–1831.

15. Wang,J., Zhuang,J., Iyer,S., Lin,X.-Y., Greven,M.C., Kim,B.-H., Moore,J., Pierce,B.G., Dong,X., Virgil,D., et al. (2013) Factorbook.org: a Wiki-based database for transcription factor-binding data generated by the ENCODE consortium. Nucleic Acids Res., 41, D171–176.

16. Kharchenko,P.V., Tolstorukov,M.Y. and Park,P.J. (2008) Design and analysis of ChIP-seq experiments for DNA-binding proteins. Nat. Biotechnol., 26, 1351–1359.

17. Levine,H.A. and Nilsen-Hamilton,M. A mathematical analysis of SELEX. Comput. Biol. Chem., 31, 11–35.

18. Tanay,A. (2006) Extensive low-affinity transcriptional interactions in the yeast genome. Genome Res., 16, 962–972.

19. Marzouk,S. El, Gahattamaneni,R., Joshi,S.R. and Scovell,W.M. (2008) The plasticity of estrogen receptor-DNA complexes: binding affinity and specificity of estrogen receptors to estrogen response element half-sites separated by variant spacers. J. Steroid Biochem. Mol. Biol., 110, 186–195.

20. Eckert,D., Buhl,S., Weber,S., Jager,R. and Schorle,H. (2005) The AP-2 family of transcription factors. Genome Biol., 6, 246.

21. Ehret,G.B., Reichenbach,P., Schindler,U., Horvath,C.M., Fritz,S., Nabholz,M. and Bucher,P. (2001) DNA binding specificity of different STAT proteins. Comparison of in vitro specificity with natural target sites. J. Biol. Chem., 276, 6675–6688.

22. Kataoka,K., Noda,M. and Nishizawa,M. (1994) Maf nuclear oncoprotein recognizes sequences related to an AP-1 site and forms heterodimers with both Fos and Jun. Mol. Cell. Biol., 14, 700–712.

23. Drew,P.D., Franzoso,G., Becker,K.G., Bours,V., Carlson,L.M., Siebenlist,U. and Ozato,K. (1995) NF kappa B and interferon regulatory factor 1 physically interact and synergistically induce major histocompatibility class I gene expression. J. Interferon Cytokine Res. Off. J. Int. Soc. Interferon Cytokine Res., 15, 1037–1045.

24. Dornan,D., Eckert,M., Wallace,M., Shimizu,H., Ramsay,E., Hupp,T.R. and Ball,K.L. (2004) Interferon regulatory factor 1 binding to p300 stimulates DNA-dependent acetylation of p53. Mol. Cell. Biol., 24, 10083–10098.

25. Abramovitch,S., Glaser,T., Ouchi,T. and Werner,H. (2003) BRCA1-Sp1 interactions in transcriptional regulation of the IGF-IR gene. FEBS Lett., 541, 149–154.

26. Chiang,B.-T., Liu,Y.-W., Chen,B.-K., Wang,J.-M. and Chang,W.-C. (2006) Direct interaction of C/EBPdelta and Sp1 at the GC-enriched promoter region synergizes the IL-10 gene transcription in mouse macrophage. J. Biomed. Sci., 13, 621–635.

27. Gartel,A.L., Ye,X., Goufman,E., Shianov,P., Hay,N., Najmabadi,F. and Tyner,A.L. (2001) Myc represses the p21(WAF1/CIP1) promoter and interacts with Sp1/Sp3. Proc. Natl. Acad. Sci. U. S. A., 98, 4510–4515.

28. Höcker,M., Raychowdhury,R., Plath,T., Wu,H., O’Connor,D.T., Wiedenmann,B., Rosewicz,S. and Wang,T.C. (1998) Sp1 and CREB mediate gastrin-dependent regulation of chromogranin A promoter activity in gastric carcinoma cells. J. Biol. Chem., 273, 34000–34007.

29. Karlseder,J., Rotheneder,H. and Wintersberger,E. (1996) Interaction of Sp1 with the growth‐ and cell cycle-regulated transcription factor E2F. Mol. Cell. Biol., 16, 1659–1667.

30. Grabowska,M.M., Elliott,A.D., DeGraff,D.J., Anderson,P.D., Anumanthan,G., Yamashita,H., Sun,Q., Friedman,D.B., Hachey,D.L., Yu,X., et al. (2014) NFI transcription factors interact with FOXA1 to regulate prostate-specific gene expression. Mol. Endocrinol. Baltim. Md, 28, 949–964.

31. Rausa,F.M., Hughes,D.E. and Costa,R.H. (2004) Stability of the hepatocyte nuclear factor 6 transcription factor requires acetylation by the CREB-binding protein coactivator. J. Biol. Chem., 279, 43070–43076.

32. Zhou,Z., Li,X., Deng,C., Ney,P.A., Huang,S. and Bungert,J. (2010) USF and NF-E2 cooperate to regulate the recruitment and activity of RNA polymerase II in the beta-globin gene locus. J. Biol. Chem., 285, 15894–15905.

33. Na,S.Y., Choi,J.E., Kim,H.J., Jhun,B.H., Lee,Y.C. and Lee,J.W. (1999) Bcl3, an IkappaB protein, stimulates activating protein-1 transactivation and cellular proliferation. J. Biol. Chem., 274, 28491–28496.

34. Baron,S., Escande,A., Albérola,G., Bystricky,K., Balaguer,P. and Richard-Foy,H. (2007) Estrogen receptor alpha and the activating protein-1 complex cooperate during insulin-like growth factor-I-induced transcriptional activation of the pS2/TFF1 gene. J. Biol. Chem., 282, 11732–11741.

35. Lee,J.S., See,R.H., Deng,T. and Shi,Y. (1996) Adenovirus E1A downregulates cJun‐ and JunB-mediated transcription by targeting their coactivator p300. Mol. Cell. Biol., 16, 4312–4326.

36. Zhang,X., Wrzeszczynska,M.H., Horvath,C.M. and Darnell,J.E. (1999) Interacting regions in Stat3 and c-Jun that participate in cooperative transcriptional activation. Mol. Cell. Biol., 19, 7138–7146.

37. Kawana,M., Lee,M.E., Quertermous,E.E. and Quertermous,T. (1995) Cooperative interaction of GATA-2 and AP1 regulates transcription of the endothelin-1 gene. Mol. Cell. Biol., 15, 4225–4231.

38. Ishiguro,A., Kassavetis,G.A. and Geiduschek,E.P. (2002) Essential roles of Bdp1, a subunit of RNA polymerase III initiation factor TFIIIB, in transcription and tRNA processing. Mol. Cell. Biol., 22, 3264–3275.

39. Xu,Z., Meng,X., Cai,Y., Koury,M.J. and Brandt,S.J. (2006) Recruitment of the SWI/SNF protein Brg1 by a multiprotein complex effects transcriptional repression in murine erythroid progenitors. Biochem. J., 399, 297–304.

40. Wadman,I.A., Osada,H., Grütz,G.G., Agulnick,A.D., Westphal,H., Forster,A. and Rabbitts,T.H. (1997) The LIM-only protein Lmo2 is a bridging molecule assembling an erythroid, DNA-binding complex which includes the TAL1, E47, GATA-1 and Ldb1/NLI proteins. EMBO J., 16, 3145–3157.

41. O’Geen,H., Lin,Y.-H., Xu,X., Echipare,L., Komashko,V.M., He,D., Frietze,S., Tanabe,O., Shi, L., Sartor,M.A., et al. (2010) Genome-wide binding of the orphan nuclear receptor TR4 suggests its general role in fundamental biological processes. BMC Genomics, 11, 689.

42. Gagliardi,A., Mullin,N.P., Ying Tan,Z., Colby,D., Kousa,A.I., Halbritter,F., Weiss,J.T., Felker,A., Bezstarosti,K., Favaro,R., et al. (2013) A direct physical interaction between Nanog and Sox2 regulates embryonic stem cell self-renewal. EMBO J., 32, 2231–2247.

43. Ambrosetti,D.C., Basilico,C. and Dailey,L. (1997) Synergistic activation of the fibroblast growth factor 4 enhancer by Sox2 and Oct-3 depends on protein-protein interactions facilitated by a specific spatial arrangement of factor binding sites. Mol. Cell. Biol., 17, 6321–6329.

44. Schwartz,C., Beck,K., Mink,S., Schmolke,M., Budde,B., Wenning,D. and Klempnauer,K.-H. (2003) Recruitment of p300 by C/EBPbeta triggers phosphorylation of p300 and modulates coactivator activity. EMBO J., 22, 882–892.

45. Zhang,Y. and Dufau,M.L. (2003) Repression of the luteinizing hormone receptor gene promoter by cross talk among EAR3/COUP-TFI, Sp1/Sp3, and TFIIB. Mol. Cell. Biol., 23, 6958–6972.

46. Formisano,L., Guida,N., Valsecchi,V., Cantile,M., Cuomo,O., Vinciguerra,A., Laudati,G., Pignataro,G., Sirabella,R., Di Renzo,G., et al. (2015) Sp3/REST/HDAd/HDAC2 Complex Represses and Sp1/HIF-1/p300 Complex Activates ncx1 Gene Transcription, in Brain Ischemia and in Ischemic Brain Preconditioning, by Epigenetic Mechanism. J. Neurosci. Off. J. Soc. Neurosci., 35, 7332–7348.

47. Carver,B.J., Plosa,E.J., Stinnett,A.M., Blackwell,T.S. and Prince,L.S. (2013) Interactions between NF-κB and SP3 connect inflammatory signaling with reduced FGF-10 expression. J. Biol. Chem., 288, 15318–15325.

48. Roder,K., Wolf,S.S., Larkin,K.J. and Schweizer,M. (1999) Interaction between the two ubiquitously expressed transcription factors NF-Y and Sp1. Gene, 234, 61–69.

49. Vallian,S., Chin,K.V. and Chang,K.S. (1998) The promyelocytic leukemia protein interacts with Sp1 and inhibits its transactivation of the epidermal growth factor receptor promoter. Mol. Cell. Biol., 18, 7147–7156.

50. Chakravarty,K., Wu,S.-Y., Chiang,C.-M., Samols,D. and Hanson,R.W. (2004) SREBP-1c and Sp1 interact to regulate transcription of the gene for phosphoenolpyruvate carboxykinase (GTP) in the liver. J. Biol. Chem., 279, 15385–15395.

51. Lim,K. and Chang,H.-I. (2010) O-GlcNAc inhibits interaction between Sp1 and sterol regulatory element binding protein 2. Biochem. Biophys. Res. Commun., 393, 314–318.

52. Biesiada,E., Hamamori,Y., Kedes,L. and Sartorelli,V. (1999) Myogenic basic helix-loop-helix proteins and Sp1 interact as components of a multiprotein transcriptional complex required for activity of the human cardiac alpha-actin promoter. Mol. Cell. Biol., 19, 2577–2584.

53. Look,D.C., Pelletier,M.R., Tidwell,R.M., Roswit,W.T. and Holtzman,M.J. (1995) Stat1 depends on transcriptional synergy with Sp1. J. Biol. Chem., 270, 30264–30267.

54. Bailey,S.D., Zhang,X., Desai,K., Aid,M., Corradin,O., Cowper-Sal Lari,R., Akhtar-Zaidi,B., Scacheri,P.C., Haibe-Kains,B. and Lupien,M. (2015) ZNF143 provides sequence specificity to secure chromatin interactions at gene promoters. Nat. Commun., 2, 6186.

55. Roopra,A., Sharling,L., Wood,I.C., Briggs,T., Bachfischer,U., Paquette,A.J. and Buckley,N.J. (2000) Transcriptional repression by neuron-restrictive silencer factor is mediated via the Sin3-histone deacetylase complex. Mol. Cell. Biol., 20, 2147–2157.

56. Kardassis,D., Falvey,E., Tsantili,P., Hadzopoulou-Cladaras,M. and Zannis,V. (2002) Direct physical interactions between HNF-4 and Sp1 mediate synergistic transactivation of the apolipoprotein CIII promoter. Biochemistry (Mosc.), 41, 1217–1228.

57. Ziegler-Heitbrock,L., Lötzerich,M., Schaefer,A., Werner,T., Frankenberger,M. and Benkhart,E. (2003) IFN-alpha induces the human IL-10 gene by recruiting both IFN regulatory factor 1 and Stat3. J. Immunol. Baltim. Md 1950, 171, 285–290.

58. Hurgin,V., Novick,D. and Rubinstein,M. (2002) The promoter of IL-18 binding protein: activation by an IFN-gamma-induced complex of IFN regulatory factor 1 and CCAAT/enhancer binding protein beta. Proc. Natl. Acad. Sci. U. S. A., 99, 16957–16962.

59. Kitabayashi,I., Yokoyama,A., Shimizu,K. and Ohki,M. (1998) Interaction and functional cooperation of the leukemia-associated factors AML1 and p300 in myeloid cell differentiation. EMBO J., 17, 2994–3004.

60. Gutierrez,S., Javed,A., Tennant,D.K., van Rees,M., Montecino,M., Stein,G.S., Stein,J.L. and Lian,J.B. (2002) CCAAT/enhancer-binding proteins (C/EBP) beta and delta activate osteocalcin gene transcription and synergize with Runx2 at the C/EBP element to regulate bone-specific expression. J. Biol. Chem., 277, 1316–1323.

61. Lively,T.N., Nguyen,T.N., Galasinski,S.K. and Goodrich,J.A. (2004) The basic leucine zipper domain of c-Jun functions in transcriptional activation through interaction with the N terminus of human TATA-binding protein-associated factor-1 (human TAF(II)250). J. Biol. Chem., 279, 26257–26265.

62. Emili,A., Greenblatt,J. and Ingles,C.J. (1994) Species-specific interaction of the glutamine-rich activation domains of Sp1 with the TATA box-binding protein. Mol. Cell. Biol., 14, 1582–1593.

63. Rossi,A., Mukerjee,R., Ferrante,P., Khalili,K., Amini,S. and Sawaya,B.E. (2006) Human immunodeficiency virus type 1 Tat prevents dephosphorylation of Sp1 by TCF-4 in astrocytes. J. Gen. Virol., 87, 1613–1623.

64. Kim,E., Yang,Z., Liu,N.-C. and Chang,C. (2005) Induction of apolipoprotein E expression by TR4 orphan nuclear receptor via 5’ proximal promoter region. Biochem. Biophys. Res. Commun., 328, 85–90.

65. Lee,J.S., Galvin,K.M. and Shi,Y. (1993) Evidence for physical interaction between the zinc-finger transcription factors YY1 and Sp1. Proc. Natl. Acad. Sci. U. S. A., 90, 6145–6149.

66. Kiryu-Seo,S., Kato,R., Ogawa,T., Nakagomi,S., Nagata,K. and Kiyama,H. (2008) Neuronal injury-inducible gene is synergistically regulated by ATF3, c-Jun, and STAT3 through the interaction with Sp1 in damaged neurons. J. Biol. Chem., 283, 6988–6996.

67. Noti,J.D. (1997) Sp3 mediates transcriptional activation of the leukocyte integrin genes CD11C and CD11B and cooperates with c-Jun to activate CD11C. J. Biol. Chem., 272, 24038–24045.

68. Lim,K. and Chang,H.-I. (2009) O-GlcNAc inhibits interaction between Sp1 and Elf-1 transcription factors. Biochem. Biophys. Res. Commun., 380, 569–574.

69. Tsai,E.Y., Falvo,J.V., Tsytsykova,A.V., Barczak,A.K., Reimold,A.M., Glimcher,L.H., Fenton,M.J., Gordon,D.C., Dunn,I.F. and Goldfeld,A.E. (2000) A lipopolysaccharide-specific enhancer complex involving Ets, Elk-1, Sp1, and CREB binding protein and p300 is recruited to the tumor necrosis factor alpha promoter in vivo. Mol. Cell. Biol., 20, 6084–6094.

70. Galvagni,F., Orlandini,M. and Oliviero,S. (2013) Role of the AP-1 transcription factor FOSL1 in endothelial cells adhesion and migration. Cell Adhes. Migr., 7, 408–411.

71. Rosmarin,A.G., Luo,M., Caprio,D.G., Shang,J. and Simkevich,C.P. (1998) Sp1 cooperates with the ets transcription factor, GABP, to activate the CD18 (beta2 leukocyte integrin) promoter. J. Biol. Chem., 273, 13097–13103.

72. Latinkić,B.V., Zeremski,M. and Lau,L.F. (1996) Elk-1 can recruit SRF to form a ternary complex upon the serum response element. Nucleic Acids Res., 24, 1345–1351.

73. Liu,X., Li,H., Rajurkar,M., Li,Q., Cotton,J.L., Ou,J., Zhu,L.J., Goel,H.L., Mercurio,A.M., Park,J.-S., et al. (2016) Tead and Ap1 Coordinate Transcription and Motility. Cell Rep., 14, 1169–1180.

74. Stewart,M.D., Choi,Y., Johnson,G.A., Yu-Lee,L., Bazer,F.W. and Spencer,T.E. (2002) Roles of Stat1, Stat2, and interferon regulatory factor-9 (IRF-9) in interferon tau regulation of IRF-1. Biol. Reprod., 66, 393–400.

75. Honda,K. and Taniguchi,T. (2006) IRFs: master regulators of signalling by Toll-like receptors and cytosolic pattern-recognition receptors. Nat. Rev. Immunol., 6, 644–658.

76. Ferreira,R., Ohneda,K., Yamamoto,M. and Philipsen,S. (2005) GATA1 function, a paradigm for transcription factors in hematopoiesis. Mol. Cell. Biol., 25, 1215–1227.

77. Woon Kim,Y., Kim,S., Geun Kim,C. and Kim,A. (2011) The distinctive roles of erythroid specific activator GATA-1 and NF-E2 in transcription of the human fetal γ-globin genes. Nucleic Acids Res., 39, 6944–6955.

78. Lee,C.S., Friedman,J.R., Fulmer,J.T. and Kaestner,K.H. (2005) The initiation of liver development is dependent on Foxa transcription factors. Nature, 435, 944–947.

79. Bonzo,J.A., Ferry,C.H., Matsubara,T., Kim,J.-H. and Gonzalez,F.J. (2012) Suppression of hepatocyte proliferation by hepatocyte nuclear factor 4α in adult mice. J. Biol. Chem., 287, 7345–7356.

80. Rodda,D.J., Chew,J.-L., Lim,L.-H., Loh,Y.-H., Wang,B., Ng,H.-H. and Robson,P. (2005) Transcriptional regulation of nanog by OCT4 and SOX2. J. Biol. Chem., 280, 24731–24737.

81. Raich,N., Clegg,C.H., Grofti,J., Roméo,P.H. and Stamatoyannopoulos,G. (1995) GATA1 and YY1 are developmental repressors of the human epsilon-globin gene. EMBO J., 14, 801–809.

82. Yardimci,G.G., Frank,C.L., Crawford,G.E. and Ohler,U. (2014) Explicit DNase sequence bias modeling enables high-resolution transcription factor footprint detection. Nucleic Acids Res., 42, 11865–11878.

83. Caminsky,N.G., Mucaki,E.J., Perri,A.M., Lu,R., Knoll,J.H.M. and Rogan,P.K. (2016) Prioritizing Variants in Complete Hereditary Breast and Ovarian Cancer (HBOC) Genes in Patients Lacking known BRCA Mutations. Hum. Mutat., 10.1002/humu.22972.

